# Phase separation of TAZ compartmentalizes the transcription machinery to promote gene expression

**DOI:** 10.1101/671230

**Authors:** Tiantian Wu, Yi Lu, Orit Gutman, Huasong Lu, Qiang Zhou, Yoav I. Henis, Kunxin Luo

**Author notes:** These authors contributed equally (Tiantian Wu, Yi Lu).

## Abstract

TAZ promotes cell proliferation, development, and tumorigenesis by regulating target gene transcription. However, how TAZ orchestrates the transcriptional responses remains poorly defined. Here we demonstrate that TAZ forms nuclear condensates via liquid-liquid phase separation to compartmentalize its DNA binding co-factor TEAD4, the transcription co-activators BRD4 and MED1 and the transcription elongation factor CDK9 for activation of gene expression. TAZ, but not its paralog YAP, forms phase-separated droplets *in vitro* and liquid-like nuclear condensates *in vivo*, and this ability is negatively regulated by Hippo signaling via LATS-mediated phosphorylation and mediated by the coiled-coil domain. Deletion of the TAZ coiled-coil domain or substitution with the YAP coiled-coil domain does not affect the interaction of TAZ with its partners, but prevents its phase separation and more importantly, its ability to induce target gene expression. Thus, our study identifies a novel mechanism for the transcriptional activation by TAZ and demonstrates for the first time that pathway-specific transcription factors also engage the phase separation mechanism for efficient transcription activation.

## Introduction

The Hippo pathway is an evolutionarily conserved signaling pathway that plays important roles in the regulation of cell proliferation, tissue homeostasis, organ size, and tumorigenesis^1-4^. At the center of this pathway is a protein kinase complex consisting of the MST1/2 (Mammalian STE20-like 1/2) and LATS1/2 (Large tumor suppressor 1/2) kinases and two accessory molecules, SAV1 and MOB1^5,6^. In response to a wide variety of signals derived from cell-cell contact, cell polarity, mechanotransduction, cellular stress and metabolism, MST1/2 is activated and in coordination with SAV1^7^, phosphorylates and activates LATS1/2. The activated LATS1/2, together with MOB1, phosphorylates the key transcription co-activators TAZ (WW domain-containing transcription regulator 1) and YAP (Yes-associated protein 1), leading to their sequestration in the cytoplasm and/or proteasomal degradation, thereby inhibiting their activity^8-12^. Once Hippo signaling is inactivated, TAZ and YAP are enriched in the nucleus, where they bind to the DNA binding co-factor TEAD and transcriptional co-activators Brd4 (Bromodomain containing 4) and MED1 (Mediator complex subunit 1)^13,14^. Through these interactions, YAP/TAZ recruits these co-activators and the transcription elongation complex to enhancers and promoters to stimulate expression of genes involved in cell proliferation, differentiation, stem cell self-renewal and carcinogenesis^13,15^. In normal tissues, well-established tissue architecture and cell-cell adhesion induce Hippo signaling to repress the activity of TAZ and YAP. In human cancer cells and tissues where proper tissue architecture is disrupted, the expression of TAZ and YAP is elevated^16^. In particular, TAZ is highly upregulated in invasive human breast cancer cell lines and in more than 20% of breast cancer tissues^17^. High levels of TAZ correlate with breast tumors of higher histological grade with increased invasiveness and an expanded cancer stem cell compartment^18^. Furthermore, overexpression of TAZ, especially the constitutively active TAZ-S89A that is resistant to LATS1/2 inhibition, in breast cancer cells promotes the expansion of cancer stem cell population and tumor invasion^18^.

TAZ and YAP are paralogs with similarities in domain structures and are subjected to inhibition by Hippo kinases in a similar manner. However, they are not functionally redundant because TAZ and YAP knockout mice show different phenotypes. For example, TAZ knockout mice are viable with defects in the kidney and lung, while YAP knockout mice are embryonic lethal with severe developmental defects^19,20^. The different functions of YAP and TAZ might be due to their differential expressions, localizations and abilities to regulate the expression of target genes in a tissue- and developmental stage-specific manner. TAZ and YAP both contain a TEAD-binding (TB) domain, WW domain(s), a coiled-coil (CC) domain, and a transcription activation (TA) domain and can bind to the same set of transcription factors including TEAD and Runx^21-23^. However, there are important differences in the sequences within these domains between TAZ and YAP that allow each of them to interact with specific transcription factors (e.g. PPARγ, TBX5, TTF-1 and Pax3 for TAZ; ErbB4 and p73 for YAP)^24^and potentially activate different target genes^25^. Although several mechanisms have been proposed to mediate transcription activation by both YAP or TAZ ^13,14,26^, the molecular mechanism underlying the functional differences between TAZ and YAP has not been well defined.

In this study, we report an important mechanism by which TAZ activates transcription through the formation of liquid-like biomolecular condensates that compartmentalize and concentrate transcription co-activators and elongation machinery. The assembly of dynamic membraneless compartments via liquid-liquid phase separation (LLPS) is essential for temporal and spatial control of numerous biochemical processes inside the cells. These membraneless compartments include many well-defined nuclear and cytoplasmic bodies such as PML (promyelocytic leukemia) bodies, Cajal bodies and stress granules, as well as subcellular structures including heterochromatin^27,28^, nuclear pore transport channels^29^, cell membrane receptor clusters^30^, and various transcription related complexes^15,31,32^. These dynamic membraneless structures, or biomolecular condensates, may serve as scaffolds to concentrate proteins that perform similar functions, or to insulate protein complexes that act in different signaling pathways to generate specificity, or to sequester proteins to facilitate or prevent inactivation. As such, they may be vital to many physiological processes, and their disruption may be associated with many pathological conditions^33^.

Proteins that tend to undergo LLPS often contain intrinsically disordered region (IDR) or are involved in weak multivalent protein-protein or protein-RNA interactions. Additional factors such as protein concentration, salt concentration, temperature, and pH also influence the ability to undergo LLPS, and various post-translational modifications can further regulate the ability of proteins to move in or out of the membraneless condensates, thereby providing efficient switch-like control for biological reactions. Recently, it has been proposed that transcriptional control may be driven by the formation of phase-separated condensates. The FET (FUS, EWS and TAF15) family of sequence-specific transcription factors, the transcription elongation factor P-TEFb as well as the coactivator proteins MED1 and BRD4 all have been demonstrated to form phase-separated condensates to activate gene expression^15,31,32,34^. Given that TAZ/YAP can interact with the transcription elongation factors and function at the super enhancers together with BRD4 and MED1^13,14^, we explored whether TAZ and YAP also form phase-separated membraneless compartments. We found that TAZ, but not YAP, undergoes LLPS through its CC domain, and these TAZ membraneless structures compartmentalize key transcriptional factors and transcription elongation machinery to facilitate gene expression. Our study has thus identified a phase separation mechanism that distinguishes TAZ from YAP to efficiently engage the transcriptional machinery for target gene expression.

## Results

### TAZ undergoes phase separation *in vitro* and *in vivo*

TAZ contains several domains important for its interactions with other cellular proteins, including the TB domain, the WW domain, the CC domain, and the TA domain^2,35^. While the crystal structure of the TA domain of TAZ in complex with TEAD4 has been reported recently^36^, the full-length TAZ protein appears to be largely unfolded^36^. Analysis of the amino acid sequence of human TAZ by the IUPred program revealed many IDRs with low complexity sequences across the entire protein (upper panels, Fig. 1a). Given that proteins containing extensive intrinsically disordered sequences or multiple modular domains tend to drive LLPS^37,38^, we asked whether TAZ could undergo phase separation. Indeed, the GFP-TAZ fusion protein purified from *E. coil* (Fig. 1a) spontaneously formed micro-sized droplets in solutions when examined under fluorescence and differential interference contrast (DIC) microscopy (Fig. 1b), and this ability was dependent on the protein concentration, salt concentration, and temperature (Figure 1b-d). GFP-TAZ formed larger and more numerous droplets at higher protein concentrations and higher salt concentrations, suggesting that the droplet formation is mediated by hydrophobic rather than electrostatic interactions. Moreover, the addition of 5% 1,6-hexanediol, a compound that putatively disrupts weak hydrophobic interactions, greatly diminished the formation of the GFP-TAZ droplets (Fig. 1e). Thus, TAZ forms phase-separated droplets in a protein concentration-, salt concentration-, and temperature-dependent manner *in vitro*.

**Fig. 1.**
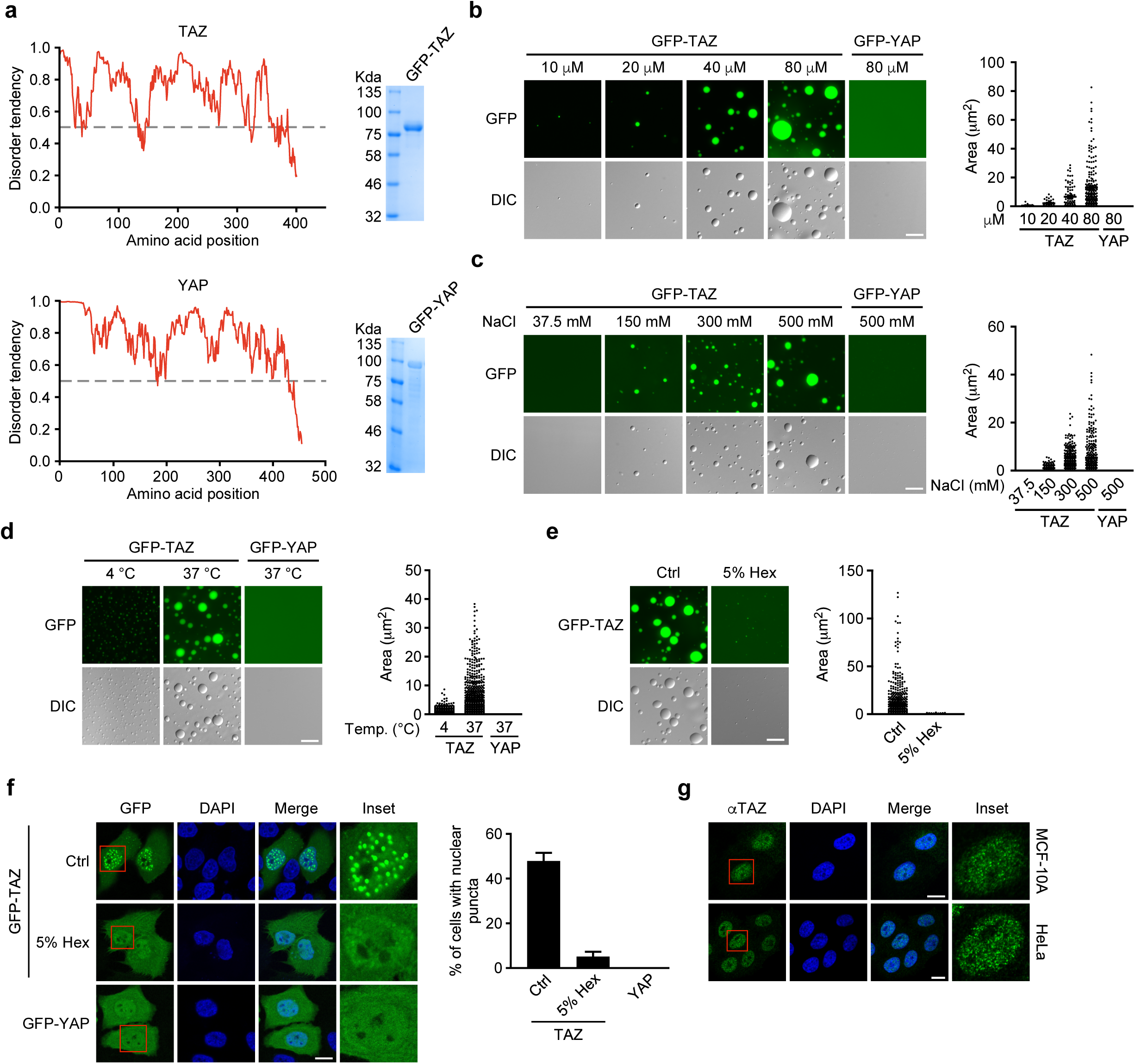
TAZ undergoes LLPS *in vitro* and *in vivo*. **a**, Graphs on the left showed the intrinsically disordered tendency of TAZ and YAP as predicted by IUPred. The program assigned scores of disordered tendency between 0 and 1 to the sequences in each protein, and a region with a score above 0.5 is considered disordered. GFP-TAZ and GFP-YAP purified from *E. coil* were analyzed by SDS-PAGE and visualized by Coomassie blue staining (Right panels). **b**, GFP-TAZ and GFP-YAP at the indicated concentrations were analyzed for their ability to form droplets at room temperature in the presence of 500 mM NaCl. Representative fluorescence and DIC images of the droplets are shown on the left, and quantification of the size and number of these droplets is shown in the graph on the right. Each dot on the graph represented a droplet. Scale bar, 10 μm. **c**, Effects of salt concentrations on droplet formation. 50 μM GFP-TAZ or GFP-YAP were subjected to droplet formation assay at room temperature in the presence of the indicated NaCl concentrations. Scale bar, 10 μm. **d**, Effects of temperature on droplet formation. 50 μM GFP-TAZ or GFP-YAP were subjected to droplet formation *in vitro* in the presence of 150 mM NaCl at 4°C or 37°C. Scale bar, 10 μm. **e**, 1,6-hexanediol disrupted droplet formation by TAZ. Droplet formation was performed with 50 μM GFP-TAZ at room temperature and 500 mM NaCl in the presence or absence of 5% 1,6-hexanediol (5% Hex). Scale bar, 10 μm. **f**, GFP-TAZ, but not GFP-YAP, formed nuclear puncta in MCF-10A cells. MCF-10A cells transfected with GFP-TAZ or GFP-YAP were treated with or without 5% Hex for 1 min and imaged. Nuclei were stained with DAPI (blue). An enlarged view of the nuclear puncta is shown in the “Inset” panel. Scale bar, 10 μm. The percentage of cells that displayed nuclear puncta is quantified in the graph to the right. **g**, Endogenous TAZ showed nuclear puncta in both MCF-10A cells and HeLa cells. TAZ was detected by immunofluorescence staining with anti-TAZ. Scale bar, 10 μm.

To test whether TAZ also undergoes LLPS *in vivo*, we first ectopically expressed GFP-TAZ in cells at a level lower than that of endogenous TAZ (Supplementary Fig. 1). Confocal microscopy revealed that GFP-TAZ formed discrete puncta in the nucleus of MCF-10A cells, which could be disrupted by treatment with 5% 1,6-hexanediol (Fig. 1f). Ectopically expressed Flag-TAZ also formed nuclear puncta, excluding the possibility that the puncta were artificially formed by the GFP tag (Supplementary Fig. 2). More importantly, the endogenous TAZ protein also exhibited nuclear puncta formation in both MCF-10A and HeLa cells (Fig. 1g). Thus, our data suggest that TAZ forms phase-separated puncta *in vivo*.

**Fig. 2.**
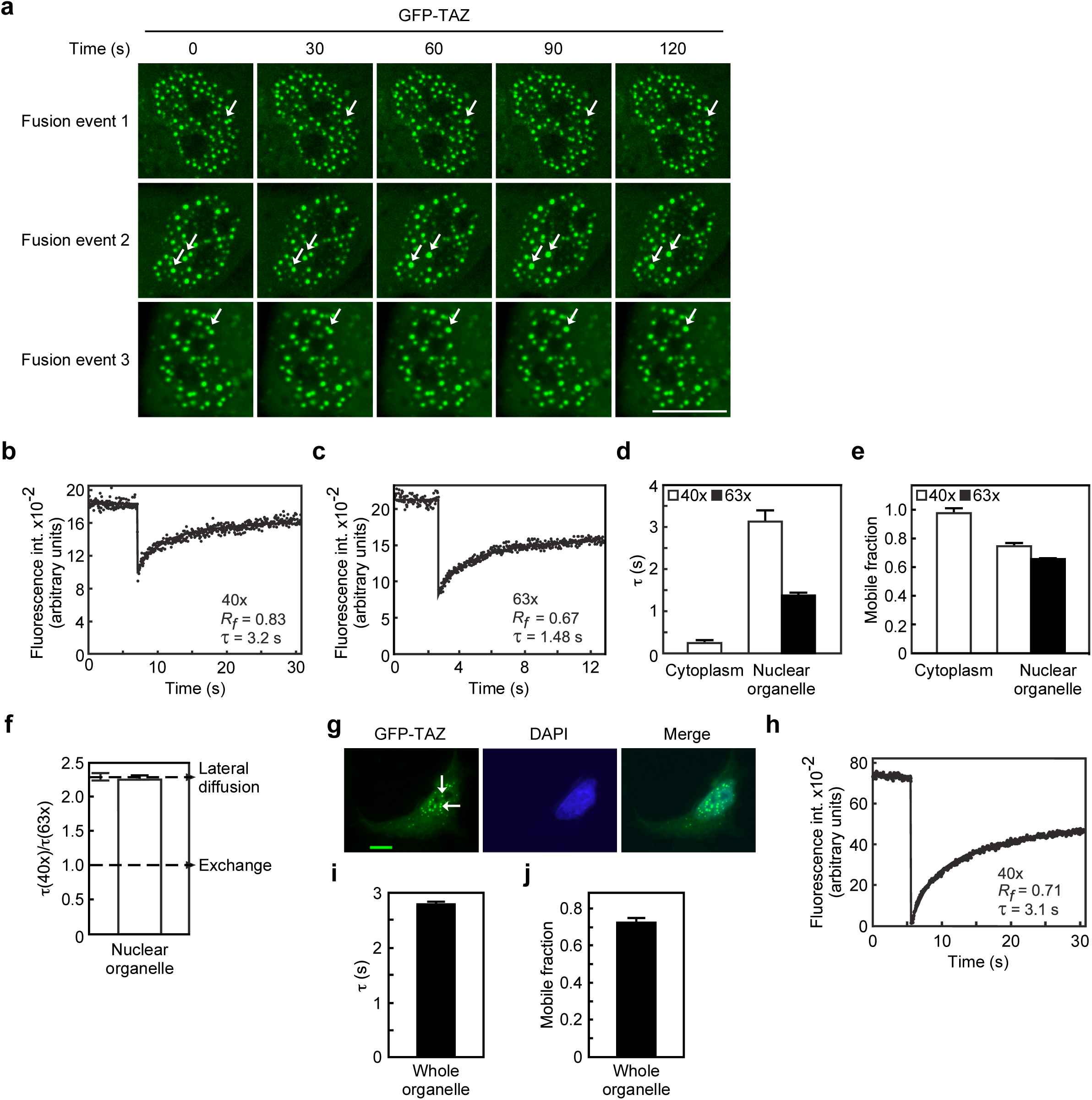
TAZ nuclear condensates display liquid-like properties. **a**, Live cell imaging of MCF-10A cells expressing GFP-TAZ. The white arrow indicates representative TAZ puncta that fused over time. **b**,**c**, Typical FRAP curves with the 40x (**b**) or 63x (**c**) objectives in spherical organelles significantly larger than the laser beam. Solid lines represent best fit by nonlinear regression to the diffusion equation (Materials and Methods); the τ and mobile fraction (*R*_*f*_) values are depicted for each panel. **d-f**, Average values and FRAP beam-size analysis. Bars are means ± SEM of 40-50 measurements on 40-50 large GFP-TAZ-containing organelles in different cells. The studies employed 40x and 63x objectives, yielding a 2.28 ± 0.06 (n=59) beam-size ratio. Thus, this τ(40x)/τ(63x) ratio is expected for FRAP by lateral diffusion (**f**, upper arrow). A τ ratio of 1 (**f**, lower arrow) indicates recovery by exchange (i.e., where the rate is determined by the chemical on and off rates). In (**f**), the SEM values of the ratios were calculated using bootstrap analysis. This analysis showed that the τ(40x)/τ(63x) ratio (2.26) of GFP-TAZ in the large organelles is similar to the 2.28 beam size ratio (*P* > 0.4), in line with FRAP time determined by diffusion. Calculating *D* from the τ values yields *D* = 0.11 ± 0.01 μm^2^/s, with *Rf* of 0.65-0.75. The τ value of GFP-TAZ in the cytoplasm (**d**), measured with the 40x objective, is over 10-fold smaller (faster diffusion), yielding *D* = 1.5 ± 0.07 μm^2^/s. **g**, A fluorescent image of GFP-TAZ organelles in the nuclei (arrow) subjected to a whole-organelle bleach with the 40x objective. The bleach time was adjusted to 150 ms to bleach the entire organelle. Bar, 10 μm. **h**, A typical FRAP curve obtained by bleaching a whole small organelle. The τ and *R*_*f*_ values are depicted. **i**,**j**, Average values of FRAP on whole organelles with approximate diameter of 1.2 μm using the 40x objective. Bars are mean ± SEM of 50 experiments on different cells (measuring one organelle in each). The τ and *R*_*f*_ values were very close to those obtained by bleaching a spot on a large organelle (compare with panels **d** and **e**). The *D* value from these experiments, based on the estimated organelle diameter, is 0.12 μm^2^/s.

### YAP differs from TAZ in its ability to undergo LLPS

YAP and TAZ are paralogs that share extensive sequence and functional similarities. Furthermore, the IUPred program also suggested extensive IDRs with low complexity sequences in YAP (lower panels, Fig. 1a). Surprisingly, under the same experimental conditions as described for TAZ above, YAP failed to form droplets *in vitro* over a wide range of protein concentrations, salt concentrations, and temperatures (Fig. 1b-d and Supplementary Fig. 3a-c). In addition, the other isoform of YAP, YAP1-2α that contains two WW domains also failed to form droplets *in vitro* (Supplementary Fig. 3d). Only in the presence of the crowding agent PEG-8000 (Polyethylene glycol 8000), did YAP form droplets (Supplementary Fig. 3e). This is consistent with a recent paper suggesting that YAP can phase separate in the presence of PEG-8000^39^. Consistent with these *in vitro* data, when ectopically expressed, GFP-YAP did not form nuclear puncta in all the cell lines tested (Fig. 1f and Supplementary Fig. 3f). Thus, YAP differs significantly from TAZ in its ability to undergo LLPS.

**Fig. 3.**
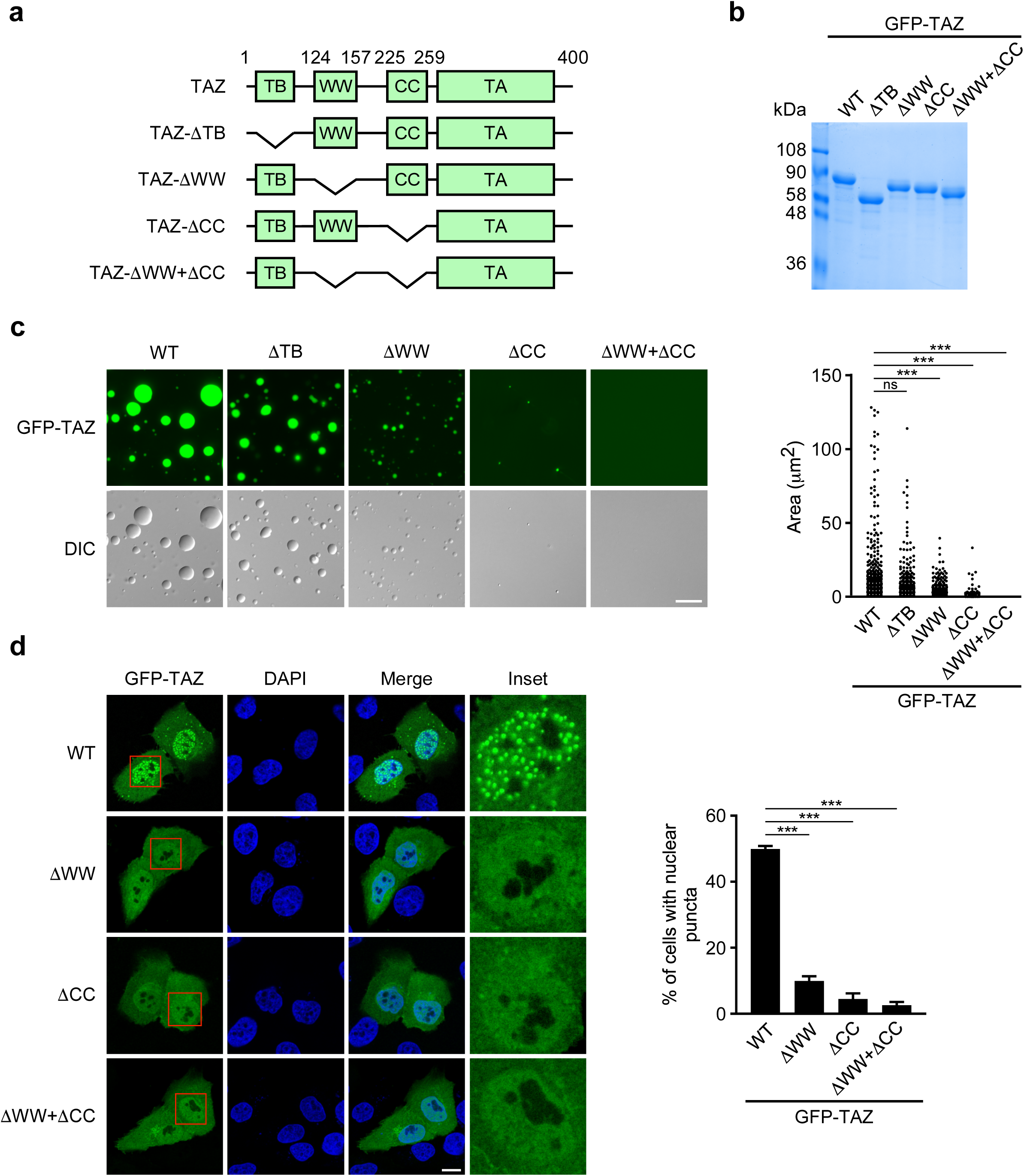
The WW domain and CC domain are required for TAZ phase separation. **a**, Domain structure of TAZ and TAZ truncations. TB: TEAD binding domain; WW: WW domain; CC: coiled-coil domain; TA: transcriptional activation domain. The numbers above indicate the position of amino acid residues. **b**, Bacterially purified GFP-TAZ, ΔTB, ΔWW, ΔCC, and ΔWW+ΔCC proteins were analyzed by SDS-PAGE and detected by Coomasssie blue staining. **c**, 50 μM WT TAZ and various mutants were subjected to droplet formation at the room temperature in the presence of 500 mM NaCl. Quantification of the droplets is on the right. Scale bar, 10 μm. ns, not significant. \*\*\**P* < 0.001. **d**, Confocal microscopy images of MCF-10A cells transfected with GFP-TAZ and various mutants (left). Scale bar, 10 μm. Quantification of the percentage of cells that displayed nuclear puncta is shown on the right. \*\*\**P* < 0.001.

### TAZ forms phase-separated puncta that exhibit liquid-like properties

Some of the criteria for defining a liquid-like phase-separated structure include a spherical shape, an ability to fuse and recovery from photo-bleaching^37,38^. We performed live cell imaging to assess whether the TAZ puncta could fuse. As shown in Fig. 2a, the TAZ nuclear condensates readily fused into larger structures over time. Next we studied the dynamics of GFP-TAZ condensates by fluorescence recovery after photobleaching (FRAP) beam-size analysis^40^, employing 63x and 40x objectives to generate two different Gaussian laser beam sizes^40^. If FRAP occurs by diffusion, τ (the characteristic fluorescence recovery time) is proportional to the bleached area (τ*_D_* = ω^2^/4*D*, where τ_*D*_ is the characteristic diffusion time, *D* the lateral diffusion coefficient, and ω is the laser beam Gaussian radius). Thus, for recovery by lateral diffusion, the ratio between the τ values obtained with the two objectives, τ(40x)/τ(63x), should equal the ratio between the bleach areas (2.28). On the other hand, when FRAP occurs by exchange with a pool of free fluorescent proteins, τ reflects the chemical relaxation time, which is independent of the bleached area, i.e. *τ*(40x)/*τ*(63x) = 1^40,41^. FRAP studies on GFP-TAZ organelles with a diameter of ∼3 μm (significantly larger than the bleach areas) yielded relatively fast recovery with τ(40x) = 3.1 s and τ(63x) = 1.4 s, with high mobile fractions (Figure 2b-e). FRAP beam-size analysis (Figure 2f) shows that the τ(40x)/τ(63x) ratio (2.23) is similar to that expected for recovery by lateral diffusion based on the ratio between the laser beam sizes (2.28). A similar value is expected for 3D diffusion in FRAP experiments involving fluorescence collection from a restricted confocal plane, since the 3D diffusion is projected into a 2D space^42^. Calculation of the lateral diffusion coefficient (*D*) from these values yields 0.11± 0.01 μm^2^/s. This value is in the same range reported for the RNA binding protein hnRNPA1 (4.2 s recovery time, with high recovery)^43^ and for an RNA helicase (2.5 s, 80% recovery, with a calculated *D* value of ∼ 0.3 μm^2^/s)^44^ in phase-separated liquid droplets in the nucleus. For comparison, GFP-TAZ in the cytoplasm displays a much faster diffusion, with *D* =1.5 μm^2^/s (Fig. 2d). Of note, bleaching whole, small GFP-TAZ organelles in the nuclei (∼ 1.2 μm diameter, using the larger beam size with the 40x objective) yielded τ of about 2.8 s with a mobile fraction above 70% (Figure 2g-j). Calculation of *D* based on the radius of the bleached organelle yielded 0.12 μm^2^/s, in line with the results obtained on large organelles, and with the reported recovery rates of RNA binding proteins upon bleaching of whole droplets^43^, suggesting that TAZ is highly dynamic, with rapid diffusion of molecules within the condensates and between them and the surrounding nuclear contents. Taken together, these results indicate that the TAZ nuclear condensates represent a separate liquid phase that is formed via LLPS.

### The CC domain is necessary for phase separation by TAZ

To identify the domains in TAZ that are required for phase separation, mutant GFP-TAZ with deletion of either the TB domain, the WW domain, or the CC domain individually, or both the WW domain and the CC domains (ΔWW+ΔCC) were purified and subjected to droplet formation *in vitro* (Fig. 3a-c). While removal of the TB domain had little effects on the ability of TAZ to form droplets, deletion of the WW domain markedly reduced, but did not eliminate, droplet formation (Fig. 3c). More strikingly, removal of the CC domain either alone or together with the WW domain abolished TAZ phase separation (Fig. 3c). These data indicate that the CC domain and to a lesser extent, the WW domain, are required for TAZ phase separation *in vitro*. Consistent with these results, deletion of the CC domain either alone or together with the WW domain significantly suppressed the ability of TAZ to form nuclear puncta *in vivo* (Fig. 3d and Supplementary Fig. 4a).

**Fig. 4.**
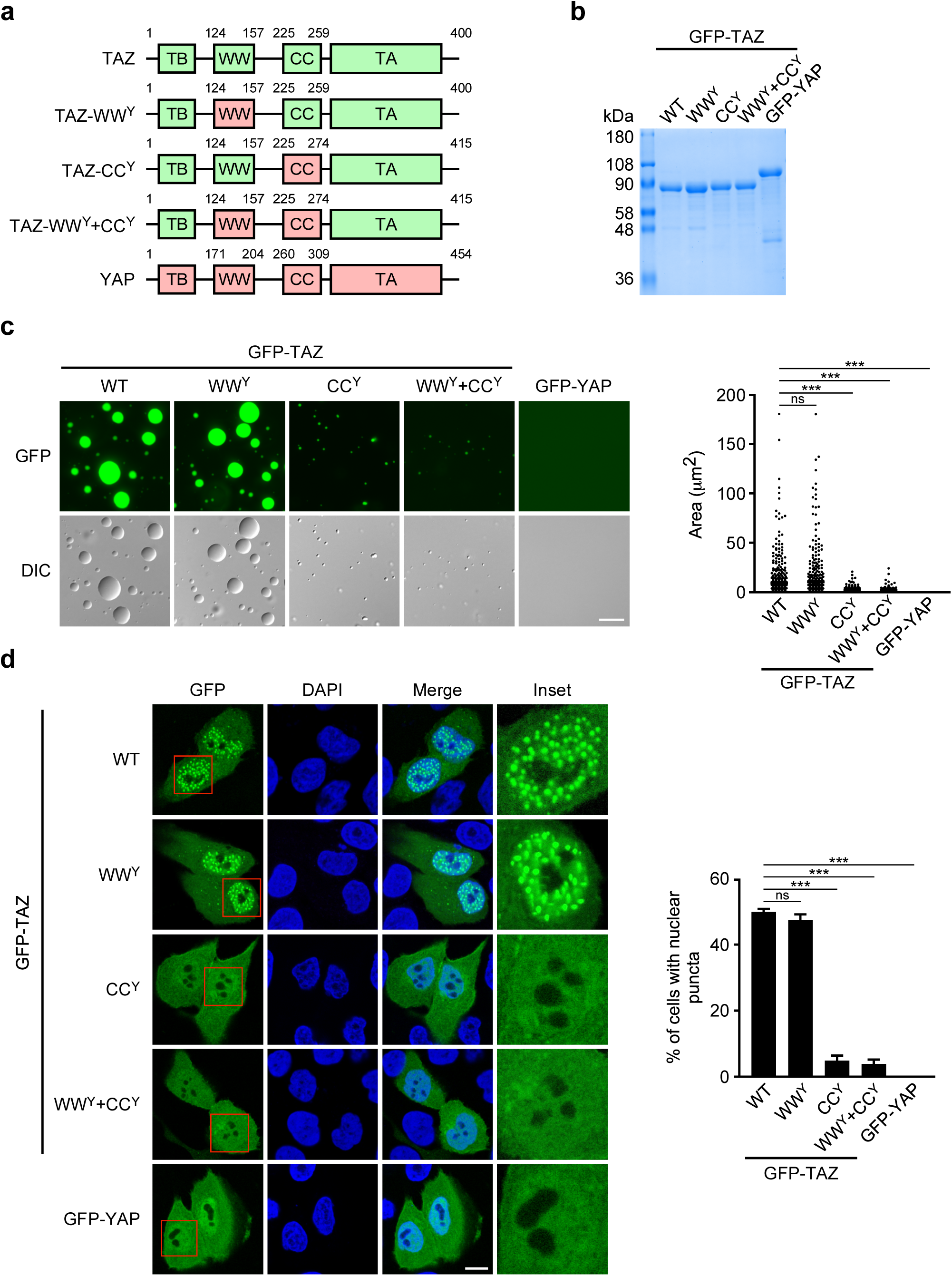
The differential ability of TAZ and YAP to undergo phase separation lies in the CC domain. **a**, Domain structure of TAZ and YAP chimera. **b**, Coomasssie blue staining of various recombinant proteins purified from *E. coil*. **c**, Droplet formation by TAZ and YAP chimera using the same condition described in Fig. 3C. Scale bar, 10 μm. Quantification of the droplets is shown on the right. \*\*\**P* < 0.001. **d**, Confocal microscopy images of MCF-10A cells transfected with various chimera as indicate. Scale bar, 10 μm. Quantification of the percentage of cells that displayed nuclear puncta is shown on the right. \*\*\**P* < 0.001.

### The CC domain distinguishes TAZ from YAP in their ability to undergo phase separation

We took advantage of the difference between YAP and TAZ in phase separation and generated TAZ/YAP chimera proteins in order to determine whether the WW and CC domains are necessary for phase separation. We swapped the WW domain and CC domain of TAZ either individually or together with that of YAP (Fig. 4a) and subjected the purified chimeric proteins to *in vitro* droplet formation assays (Fig. 4b-c). While full-length TAZ and chimeric TAZ containing the YAP WW domain (WW^Y^) readily formed droplets, TAZ chimera containing the YAP CC domain (CC^Y^) or the YAP WW and CC domains (WW^Y^+CC^Y^) failed to do so (Fig. 4c). Consistent with the results from the *in vitro* assays, TAZ chimera containing either the YAP CC domain alone or the YAP CC and WW domains did not display nuclear puncta *in vivo*, but that containing the YAP WW domain exhibited similar nuclear puncta pattern as wild-type TAZ (Fig. 4d and Supplementary Fig. 4b). Thus, it appears that the difference in the CC domain sequences between TAZ and YAP determined their differential ability to undergo phase separation.

### TAZ phase separation is negatively regulated by Hippo signaling via phosphorylation by LATS

TAZ is an important downstream effector of the Hippo signaling pathway, and its localization and activity can be negatively regulated by Hippo signaling. We next asked whether the ability of TAZ to undergo LLPS could be regulated by Hippo signaling. To this end, we subjected the cells to serum stimulation or alteration in cell density, two processes known to regulate the Hippo kinases. Upon serum starvation, ectopically expressed or endogenous TAZ was diffusely localized in the cytoplasm, as published previously^45,46^. The addition of serum led to the translocation of TAZ to the nucleus and in particular, to the nuclear puncta (Fig. 5a,b). Similarly, the formation of TAZ nuclear condensates was also regulated by cell density. At the high cell density, TAZ was restricted to the cytosol in a diffuse manner or degraded as a result of inactivation by the Hippo kinases^2,45^. In contrast, at the low cell density, the stabilized TAZ translocated into the nucleus and clearly formed nuclear condensates (Fig. 5a,b). Thus, activated Hippo signaling inhibits LLPS of TAZ.

**Fig. 5.**
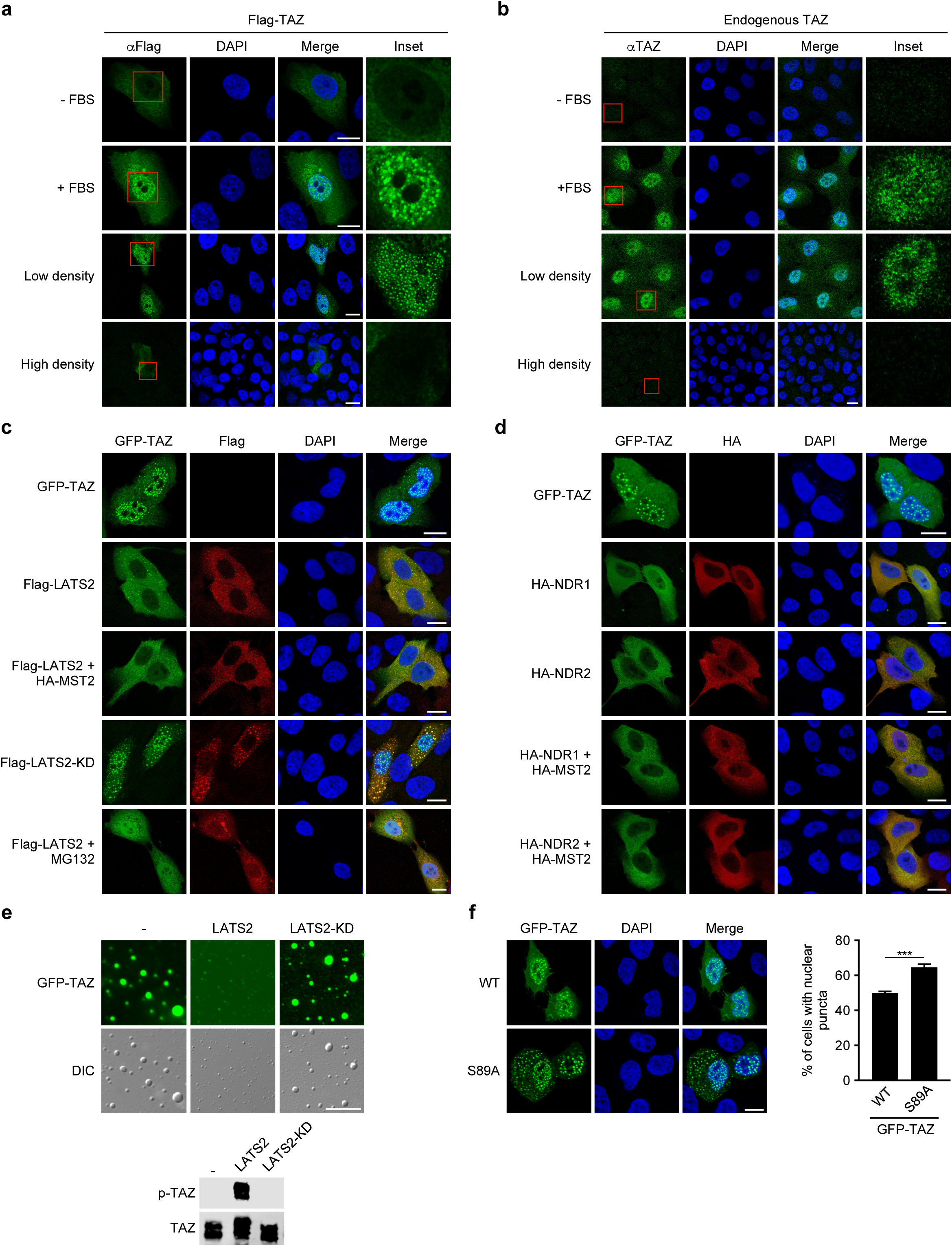
Hippo signaling negatively regulates TAZ phase separation through LATS2. MCF-10A cells transfected with Flag-TAZ (**a**) or not (**b**) were serum-starved for 16 h, followed by treatment with 10% FBS; or cultured at low density or high density. Localization of Flag-TAZ (**a**) or endogenous TAZ (**b**) was detected by immunofluorescence with anti-Flag (green) or anti-TAZ (green) antibodies. Scale bars, 10 µm. **c**, GFP-TAZ (Green) was co-transfected with WT LATS2, either alone or together with HA-MST2, or with the kinase inactive LATS2-KD in the absence of presence of 40 µM MG132 for 6 h. LATS2 localization was detected by immunofluorescence with anti-Flag (Red). Scale bars, 10 µm. **d**, GFP-TAZ (Green) was co-transfected with WT Flag-NDR1 or NDR2, either alone or together with HA-MST2. NDR1/2 location was detected by immunofluorescence with anti-Flag (Red). Scale bars, 10 µm. **e**, *In vitro* phosphorylation and droplet formation. GFP-TAZ was phosphorylated in an *in vitro* kinase assay by WT or kinase inactive LATS2 prepared from transfected 293T cells and subjected to the droplet formation assay. Phosphorylation of TAZ was detected by western blotting using antibodies specific for phosphor-TAZ (lower panel). Representative fluorescence and DIC images of the droplets are shown on the top. Scale bars, 10 µm. **f**, Confocal images of MCF-10A cells transfected with GFP-TAZ or GFP-TAZ-S89A. Scale bars, 10 µm. Quantification of the percentage of cells that displayed nuclear puncta is shown on the right. \*\*\**P* < 0.001.

Hippo signaling activates the LATS kinases that phosphorylate TAZ at Ser 89, leading to its cytoplasmic localization and subsequent proteasomal degradation^22^. We next investigated whether LATS2 could directly affect the phase separation of TAZ through phosphorylation. Ectopic expression of LATS2 either alone or together with MST2 inhibited the formation of nuclear puncta by TAZ, whereas the kinase-inactive LATS2-KD failed to block this process (Fig. 5c), suggesting that phosphorylation of TAZ by LATS2 prevented phase separation. Since phosphorylation of TAZ by LATS2 results in its degradation and exclusion from the nucleus, which could account for the lack of nuclear puncta formation, we next blocked its degradation by treating the cells with the proteasome inhibitor MG132. The MG132-treated cells retained most TAZ in the nucleus, but the phosphorylated TAZ did not form nuclear puncta (Fig. 5c). In addition to LATS2, NDR1/2, two other Ser/Thr kinases of the NDR/LATS family, also phosphorylate TAZ^47^. Similar to LATS2, ectopic expression of NDR1 or NDR2 either alone or together with MST2 significantly impaired the ability of TAZ to form nuclear condensates (Fig. 5d). To unequivocally confirm that phosphorylation of TAZ by LATS2 is responsible for the inhibition of phase separation, we performed an *in vitro* kinase assay using LATS2 to phosphorylate purified GFP-TAZ, and examined the ability of this phosphorylated TAZ to form droplets *in vitro*. As shown in Fig. 5e, phosphorylated TAZ exhibited a greatly reduced ability to form droplets *in vitro*. Moreover, the TAZ-S89A mutant, which is resistant to LATS2 phosphorylation, displayed moderately enhanced condensates formation when compared to wild-type TAZ (Fig. 5f). Taken together, these data indicate that the ability of TAZ to undergo LLPS can be inhibited by Hippo signaling via LATS/NDR kinases-mediated phosphorylation.

### Phase-separated TAZ can compartmentalize TEAD and other transcription co-factors

To determine the potential functions of TAZ phase separation, we investigated what molecules co-segregated with TAZ condensates in the nucleus by immunofluorescence using antibodies specific for known markers of nuclear organelles or condensates. Our initial survey indicated that TAZ nuclear condensates did not co-localize with markers of commonly observed nuclear bodies, such as PML (PML bodies), Fibrillarin (nucleolus), or Coilin (Cajal bodies) (Supplementary Fig. 5). We next asked whether the TAZ nuclear puncta were enriched with its DNA binding co-factors TEAD^10,48,49^ and other known transcriptional co-activators. As shown in Fig. 6a, while TEAD4 alone was evenly distributed in the nucleus, in the presence of ectopically expressed GFP-TAZ but not GFP-TAZ-S51A deficient in TEAD binding, TEAD4 was directed into the nuclear condensates and co-localized with TAZ. Deletion of the WW and CC domains of TAZ (ΔWW+ΔCC) not only disrupted the formation of the TAZ nuclear puncta but also prevented co-localization of TEAD4 in these puncta, even though this TAZ mutant could still bind to TEAD4 (Fig. 6c). This suggests that the enrichment of TEAD4 in the nuclear puncta was dependent upon TAZ. Consistently, *in vitro* droplet formation assay further confirmed that TEAD4 alone did not form droplets, but was recruited to these droplets by TAZ (Fig. 6b). Again, deletion of the WW and CC domains prevented droplet formation *in vitro* by both TAZ and TEAD4. These data suggest that TAZ interacts with TEAD4 and recruits TEAD4 to the liquid droplets.

**Fig. 6.**
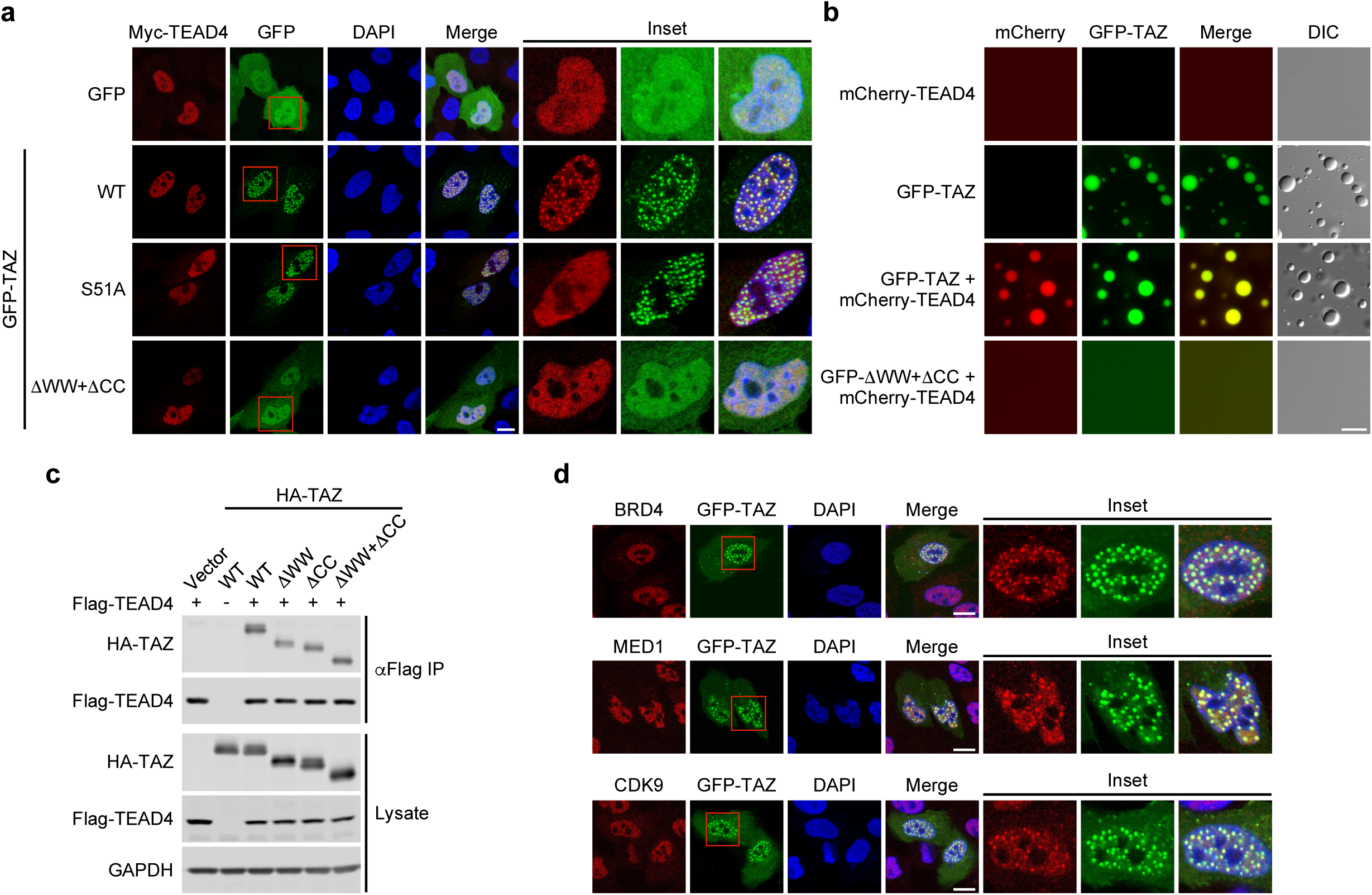
TAZ compartmentalizes TEAD and other transcriptional factors to the nuclear puncta. **a**, Myc-TEAD4 was co-transfected into MCF-10A cells together with GFP vector, WT GFP-TAZ, or S51A or ΔWW+ΔCC mutant. TAZ localization was monitored by GFP, and TEAD4 by immunofluorescence staining with anti-Myc (Red). Scale bar, 10 μm. **b**, *In vitro* droplet formation assay. 50 μM mCherry-TEAD4 either alone or mixed together with 50 μM WT GFP-TAZ or ΔWW+ΔCC were subjected to the droplet formation assay using the same condition as described in Fig. 3C. Scale bar, 10 μm. **c**, The ability of HA-tagged WT or mutant TAZ to interact with Flag-TEAD4 was examined by a co-immunoprecipitation (co-IP) assay using the anti-Flag antibody in the immunoprecipitation (IP), followed by western blotting with anti-HA (upper). The abundance of these proteins in the cell lysates was assessed by western blotting (lower). **d**, Co-localization of BRD4, MED1 or CDK9 with GFP-TAZ in the nuclear puncta *in vivo*. Localization of endogenous BRD4, MED1 and CDK9 was detected by indirect immunofluorescence (Red). Scale bar, 10 μm.

In addition to TEAD4, the TAZ nuclear condensates were also enriched with component of the transcriptional elongation machinery, CDK9, and super-enhancer markers BRD4 and MED1 (Fig. 6d), further supporting the idea that the TAZ population in these nuclear puncta is transcriptionally active. Taken together, these data support the model that TAZ forms nuclear condensates that are enriched with key transcription factors and machinery to facilitate transcription of target genes. This is consistent with the recent report that TAZ binds to BRD4 and is associated with selected super-enhancers, recruiting BRD4 to chromatin to regulate TAZ target gene expression^14^.

### Phase separation promotes transcriptional activation by TAZ

We next examined whether the ability of TAZ to undergo phase separation is required for its transcriptional activity. We took advantage of the TAZ ΔCC mutant and TAZ CC^Y^chimera that do not form nuclear puncta, but are still able to bind to its DNA binding co-factor TEAD4 and its regulator LATS2 (Fig. 6c and Supplementary Fig. 6). In a TAZ-dependent luciferase reporter assay, deletion of the WW domain alone had little effect on transcription, while deletion of the CC domain either individually or together with the WW domain significantly decreased its transcriptional activity (Fig. 7a). Consistent with this result, expression of two endogenous TAZ target genes, *CTGF* and *CYR61* was induced to a less extent by TAZ mutant lacking the CC domain either alone or together with the WW domain (Fig. 7b). Thus, the transcriptional activity of TAZ correlated with its ability to undergo LLPS.

**Fig. 7.**
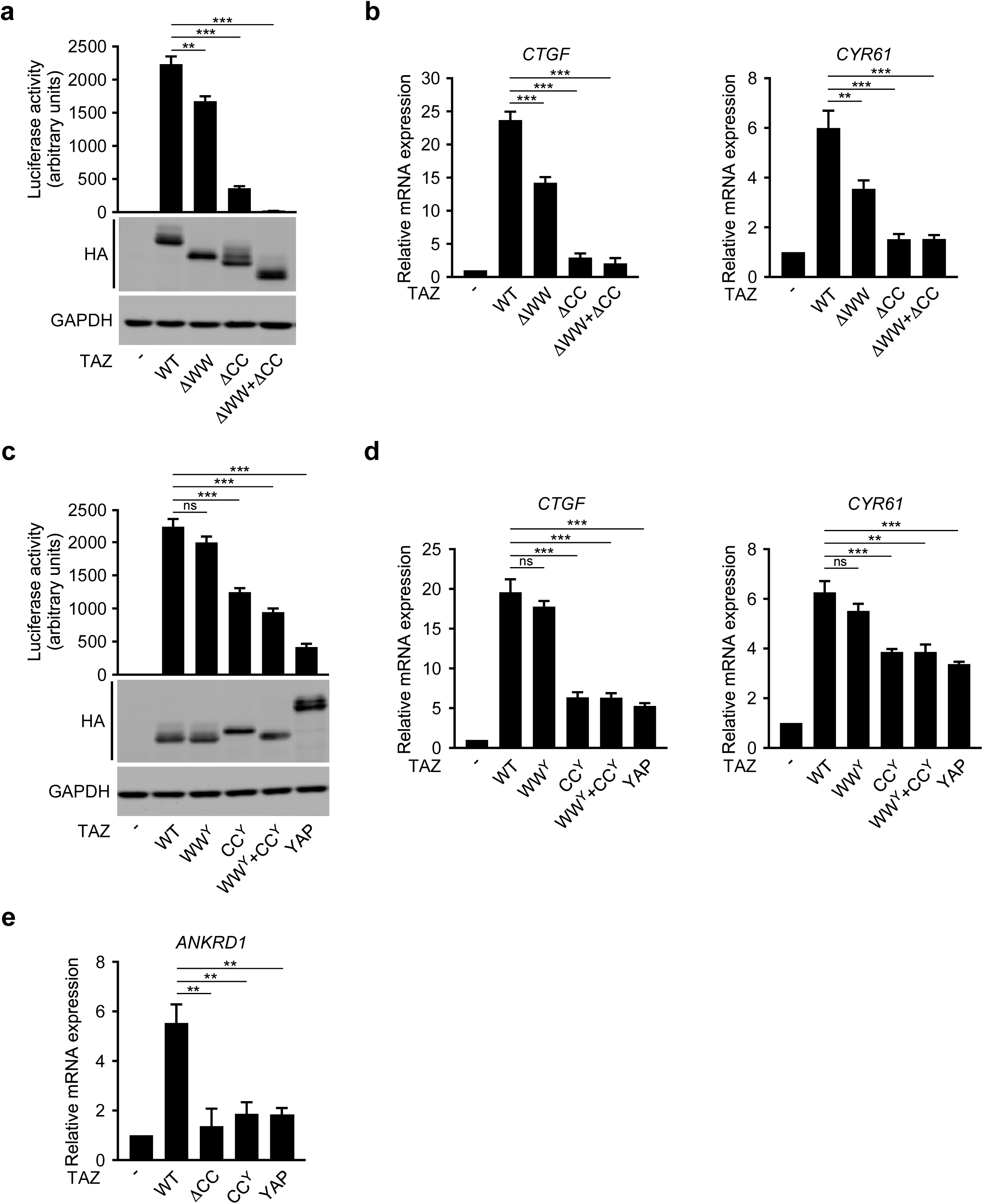
Phase separation of TAZ promotes transcription. **a**,**c**, The TAZ-dependent luciferase activity was measured in 293T cells expressing 8xGT-IIC-δ51LucII and various TAZ mutants (**a**) or TAZ/YAP chimera (**c**). Each data point represents the mean ± SEM from duplicate samples. **b**,**d**,**e**, qRT-PCR analysis of *CTGF, CYR61*, and *ANKRD1* mRNA expression in 293T cells transfected with various TAZ mutants (**b**,**e**) or TAZ/YAP chimera (**d**,**e**).

Although both TAZ and YAP are transcription activators, under the same condition, YAP exhibited a lower transcription activity than TAZ (Fig. 7c-d)^36^. Interestingly, while TAZ-WW^Y^ had a similar transcription activity as TAZ, TAZ-CC^Y^ and TAZ-WW^Y^+CC^Y^ displayed reduced transcriptional activation at a level similar to that of YAP in both the luciferase reporter assay and in expression of endogenous target genes (Fig. 7c-d). Similarly, expression of *ANKRD1*, a gene known to be sensitive to TAZ regulation in many cell types^46^, was also impaired by the deletion or substitution of TAZ CC domain (Fig. 7e). Taken together, these data support a correlation between the ability of TAZ to undergo phase separation and its transcription activity. Since YAP and TAZ-CC^Y^ are capable of activating transcription, albeit at a lower level, the TAZ phase separation is probably not essential for the basal transcription activity, but is crucial for optimal transcription.

## Discussion

The Hippo pathway transcription effectors TAZ and YAP have been shown to activate the expression of many cellular genes in response to a wide variety of signals derived from cell-cell contact, cell polarity, mechano-transduction, cellular stress and metabolism^50-53^. How a broad spectrum transcriptional co-activator like TAZ or YAP orchestrates such a diverse array of signals to generate specific downstream outcomes is an important but unanswered question. Here we show that phase separation can serve as an important mechanism that allows TAZ to activate transcription in an efficient and specific manner. TAZ, but not YAP, forms liquid-like droplets both *in vitro* and *in vivo*, and these phase-separated particles serve as hubs for TAZ to recruit and compartmentalize its partner DNA binding cofactor TEAD4 and transcription co-activators and core machinery including BRD4, MED1 and CDK9 to activate target genes transcription. Mutant TAZ (ΔCC) defective in phase separation but capable of binding to its partners TEAD4 or LATS2 fails to activate target gene expression. Consistently, the ability of TAZ to phase separate is negatively regulated by Hippo signaling via phosphorylation by LATS/NDR kinases. Signals that activate Hippo signaling such as serum starvation or high cell density all prevent TAZ nuclear phase separation. Thus, our study has identified a novel mechanism by which TAZ efficiently engages the transcriptional machinery to stimulate gene expression.

YAP differs from TAZ in its ability to phase separate. Under the condition where TAZ readily undergoes phase separation, YAP fails to form droplets either *in vitro* or *in vivo*. Only in the presence of the crowding agent PEG-8000, did YAP form droplets. The ability of a given protein to undergo phase separation is often influenced by the presence of intrinsically disordered regions or CC domains that mediate oligomerization or interaction with other proteins necessary for assembly of liquid-like condensates^54,55^. For example, the formation of germ granules and PML bodies is initiated by self-association of the CC domains of the PGL-1/3 and PML proteins, respectively^56^. In these cases, modular domains mediate specific interactions to form structures that serve as scaffolds for further assembly of components of the membrane-less organelles. TAZ and YAP both contain large stretches of intrinsically disordered regions and a CC domain. We found that the differential ability of TAZ and YAP to undergo phase separation is determined by the CC domain. Deleting the TAZ CC domain or substituting it with the YAP CC domain significantly impaired the ability of TAZ to undergo phase separation and more importantly, to activate target gene expression. The CC domain could facilitate TAZ phase separation by mediating potential TAZ oligomerization and/or multivalent interactions between TAZ and other cellular proteins. These oligomerization/multivalent protein interactions likely involve hydrophobic interactions as TAZ phase separation is promoted by increasing salt concentrations and temperature, which are conditions favoring hydrophobic interactions. The TAZ CC domain has been reported previously to mediate interaction with the Smad2/3-4 complex to promote their nuclear translocation^57^. However, its role in transcriptional activation has not been previously defined. Our study thus demonstrates for the first time a critical role of the CC domain in transcriptional activation through inducing phase separation.

Although TAZ and YAP share a high level of sequence similarities, and both are regulated by Hippo signaling, they exhibit different functions *in vivo*^19,20^. Supporting these functional differences, gene expression microarray and RNA-sequencing analyses have shown that although many genes can be activated by both TAZ and YAP, there are also a significant subset of genes that are differentially induced by either TAZ or YAP, often in a cell- or tissue-specific manner^25,58^. How TAZ and YAP achieve this functional specificity at the transcription level is not well understood. Kaan et al. suggest that, unlike the YAP-TEAD dimer, TAZ-TEAD can form a hetero-tetramer, and this differential structural feature may affect DNA target selectivity and transcription of some target genes^36^. Our results suggest that phase separation may allow TAZ to compartmentalize and concentrate the transcription co-activators and general transcription machinery in dynamic membraneless domains, thereby leading to more efficient transcription reactions. This is consistent with the finding that TAZ may be a more effective transcription activator than YAP under overexpression conditions^36^. The compartmentalization of the TAZ-initiated transcription complex may also physically separate the TAZ-specific signaling pathways from those specific for YAP and thus provide pathway specificity. Taken together, our results provide a novel mechanistic basis for the differential transcriptional activities of TAZ and YAP.

LLPS is emerging as a key mechanism that is critical for transcriptional regulation. General transcription elongation factor P-TEFb, transcription initiation factor TAF15 and FUS^59,60^as well as stem-cell specific transcription factors OCT4, MYC and SOX2^15^, have all been shown to undergo phase separation to cluster in discrete membraneless condensates that function as hubs to allow efficient and dynamic regulation of transcription and RNA processing. More recently, bromo-domain transcription factor BRD4 and MED1 have been reported to form phase-separated condensates at super enhancers that compartmentalize and concentrate the transcription apparatus for gene expression^31^. Here we provide the first evidence that the signaling pathway-specific transcription co-activator TAZ employs the phase separation mechanism via multivalent protein-protein interactions through its CC domain to regulate downstream gene expression. In this case, the formation of phase-separated nuclear speckles by TAZ allows recruitment and enrichment of its DNA-binding co-factor TEAD4, transcription co-activators BRD4 and MED1 as well as the general elongation factor P-TEFb in one compartment. This arrangement allows TAZ to efficiently orchestrate the entire transcriptional machinery to regulate the expression of target genes. Indeed, mutations in TAZ that disrupt its phase separation but still retain interactions with its partners and regulators TEAD4 and LATS1/2 showed greatly diminished transcriptional activity. Compartmentalization of TAZ in transcriptionally active condensates also spatially separates it from its upstream regulator LATS1/2, effectively insulating this population from inactivation. As TAZ is a critical regulator of cell proliferation, survival, differentiation, and transformation^14,61^ and its up-regulation in human cancers can promote transcriptional addiction, understanding the role of phase separation in its mechanism of action may provide new therapeutic targets for human cancer.

## Methods

### Plasmids, antibodies, and reagents

The GFP-TAZ and GFP-YAP constructs were generated by PCR and sub-cloned into the pEGFP-C1 vector (Clontech) or pGFP-2xStrep vector, kindly provided by Dr. Qiang Zhou (University of California, Berkeley). Mutant GFP-TAZ containing various truncations and mutations in the TAZ molecule were generated by PCR and similarly cloned into the above vectors. The chimeric GFP-TAZ molecules containing the substituted YAP WW (WW^Y^: aa 171-204), CC (CC^Y^: aa 260-309), or both WW and CC domains (WW^Y^+CC^Y^) were generated based on GFP-TAZ by PCR. The mCherry-TEAD4 construct was generated by PCR and sub-cloned into the pHis-mCherry vector provided by Dr. Qiang Zhou (University of California, Berkeley). cDNAs of TAZ, YAP, LATS2, LATS2-KD, TEAD4, and MST2 were kindly provided by Dr. Kun-Liang Guan (University of California, San Diego) and Dr. Alain Mauviel (Curie Institute, Orsay, France).

The following antibodies and reagents were purchased from commercial sources: TAZ (BD Pharmingen™, 560235); GAPDH (Santa Cruz, FL-335); GFP (Thermo Fisher, GF28R); Myc (Cell Signaling Technology, 9B11); MED1 (Santa Cruz, M-255); PML (Santa Cruz, PG-M3); Coilin (Santa Cruz, F-7); Fibrillarin (Santa Cruz, G-8); Flag (Sigma, F3165); MG-132 (Selleck), 1,6-hexanediol (Sigma), and PEG-8000 (Sigma).

Antibodies against CDK9, BRD4, and HA were generated and kindly provided by Dr. Qiang Zhou (University of California, Berkeley) as described previously (Lu et al., 2018).

### Protein expression and purification

Plasmids containing Strep-GFP- or His-mCherry-tagged genes were transformed into *E. coli* BL21 cells. After induction with IPTG, bacteria lysates in buffer (50 mM Tris-HCl pH7.5, 500 mM NaCl, 1 mM DTT, 1% Triton X-100) were sonicated, and the Strep-GFP-fusion proteins were purified using the Strep-Tactin Superflow beads (IBA). The His-mCherry-fusion proteins were purified using a Ni-NTA column (Thermo Fisher Scientific). The eluted proteins were dialyzed in 1 L dialyzed buffer (20 mM Tris-HCl pH 7.5, 37.5 mM NaCl, and 1mM DTT) overnight at 4°C and concentrated with Amicon ultra centrifugal filters (Millipore).

### Droplet formation assay

Purified proteins were diluted to varying concentrations in buffer containing 20 mM Tris-HCl pH 7.5 and 1 mM DTT with the indicated salt concentrations. 5 μL of the protein solution was loaded onto a glass slide, covered with a coverslip, and imaged with AxioObserver Z1 inverted microscope (Zeiss). The sizes of the droplets in three 166×124 μm^2^ fields were quantified by ImageJ.

### Cell culture and transfection

293T and HeLa cells were cultured in DMEM (Invitrogen) containing 10% FBS (HyClone) and 50 μg/mL penicillin/streptomycin (Pen/Strep). MCF10A cells were cultured in DMEM/F12 (Invitrogen) supplemented with 5% horse serum (Invitrogen), 20 ng/mL EGF, 0.5 μg/ml hydrocortisone, 10 μg/ml insulin, 100 ng/ml cholera toxin, and 50 μg/mL Pen/Strep. All cell lines were authenticated at UC Berkeley Cell Culture Facility by single nucleotide polymorphism testing and confirmed as mycoplasma negative.

Transfection of cells was performed using Lipofectamine 3000 (Thermo Fisher Scientific) according to manufacturer’s instruction.

### Immunofluorescence staining and live cell imaging

Cells were seeded on glass coverslips, fixed with 4% paraformaldehyde-PBS for 15 min, blocked in buffer containing 5% FBS and 0.3% Triton X-100 in PBS for 1 h, and incubated with primary antibodies overnight at 4°C. After washes, cells were incubated with Alexa Fluor 488- or 555-conjugated secondary antibodies (Thermo Fisher Scientific) for 1 h at room temperature. Immunofluorescence was detected using Zeiss LSM 710 confocal microscopy.

Live cell imaging was performed as previously described^32^. Briefly, MCF-10A cells transfected with GFP-TAZ construct were seeded on LabTek chambered slides (Nunc, Denmark) and examined under a Nikon Spinning Disk confocal microscope. During image acquisition, cells were incubated in an equilibrated observation chamber at 37°C and with 5% CO2. Images were acquired at 30-second intervals and were loaded in ImageJ to identify fusion events.

### FRAP and FRAP beam-size analysis

HeLa cells grown on glass coverslips in 6-well plates were transfected with 2 μg/well of GFP-TAZ and subjected to Quantitative FRAP studies 24 h post transfection as described previously^40,62^. Measurements were performed at 22 °C in Hank’s balanced salt solution supplemented with 20 mM HEPES, pH 7.2 (HBSS/HEPES). An argon ion laser beam (Innova 70C; Coherent) was focused through a fluorescence microscope (AxioImager.D1, Carl Zeiss MicroImaging) to a spot with a Gaussian radius (ω) of 0.77 ± 0.03 μm (Plan apochromat 63x/1.4 NA oil-immersion objective) or 1.17 ± 0.05 μm (C apochromat 40x/1.2 NA water-immersion objective). The ratio between the bleach areas [ω^2^(40x)/ω^2^(63x)] was 2.28 ± 0.06 (n = 59; SEM calculated using bootstrap analysis as described below). After a brief measurement at monitoring intensity (488 nm, 1 μW), a 5 mW pulse (5-10 ms) bleached 60–75% of the fluorescence in the illuminated region, and recovery was followed by the monitoring beam. The characteristic fluorescence recovery time (τ) and mobile fraction (*R*_*f*_) were extracted by nonlinear regression analysis, fitting to a lateral diffusion process^40^. The significance of differences between τ values measured with the same beam size was evaluated by Student’s *t*-test. To compare ratio measurements [τ(40x)/τ(63x) and ω^2^(40x)/ω^2^(63x)], we employed bootstrap analysis, which is preferable for comparison between ratios^63^, as described earlier^62^, using 1,000 bootstrap samples.

### Immunoprecipitation and immunoblotting

Immunoprecipitation and Immunoblotting were performed as previously described^64^. Briefly, cells were lysed in lysis buffer (50 mM HEPES at pH 7.5, 150 mM NaCl, 1 mM EDTA, 1% NP-40, 10 mM pyrophosphate, 10 mM glycerophosphate, 50 mM NaF, 1.5 mM Na_3_VO_4_, protease inhibitor cocktail [Roche], 1 mM PMSF), and clarified cell lysates were subjected to immunoprecipitation with anti-Flag M2 agarose beads (Sigma).

### *In vitro* kinase assay

Flag-LATS2 or Flag-LATS2-KD was purified from transfected 293T cells and eluted as described previously^64^, and incubated with GFP-TAZ immobilized on the Strep-Tactin Superflow beads in kinase buffer (10 mM MgCl_2_, 50 Mm NaCl, 1 mM DTT, 50 mM HEPES at pH 7.4) containing 5mM ATP at room temperature for 12 h. After washing, the phosphorylated GFP-TAZ were eluted from the beads with 10 μL elution buffer (20 mM Tris-HCl pH 7.5, 37.5 mM NaCl, 1 mM DTT and 1 mM desthiobiotin).

### Luciferase assay

A total of 2.5 μg DNA (including 50 ng of 8xGT-IIC-δ51LucII Luciferase reporter construct and the indicated plasmids) was transiently transfected into 293T cells by Lipofectamine 2000. The luciferase activity was measured at 36 h after transfection as described previously^64,65^.

### RNA extraction, reverse transcription, and real-time PCR

Total RNA was extracted using TRIzol Reagent (Ambion). 1 μg RNA was reverse-transcribed using the SuperScript™ III First-Strand Synthesis System (Thermo Fisher Scientific). The resulted cDNA was subjected to quantitative real-time PCR (qRT-PCR) using the DyNAmo HS SYBR Green qPCR Kits (Thermo Fisher Scientific) and the Bio-Rad real-time PCR system (Bio-Rad), with *β-actin* gene as a control. Primers used are: *β-actin*, Forward: GCCGACAGGATGCAGAAGGAGATCA, Reverse: AAGCATTTGCGGTGGACGATGGA; *CTGF*, Forward: CCAATGACAACGCCTCCTG, Reverse: TGGTGCAGCCAGAAAGCTC; *CYR61*, Forward: AGCCTCGCATCCTATACAACC, Reverse: TTCTTTCACAAGGCGGCACTC; *ANKRD1*, Forward: CACTTCTAGCCCACCCTGTGA, Reverse: CCACAGGTTCCGTAATGATTT

### Statistical analysis

All data were derived from at least three independent experiments and are presented as means ± SEM. Comparisons among groups were performed by one-way ANOVA (Krusk-Wallis test) or Student’s *t*-test with Prism5. FRAP beam-size ratio measurements employed bootstrap analysis, which is preferable for comparison between ratios^63^, as described earlier^62^, using 1,000 bootstrap samples. *P* values are shown when relevant (\**P* < 0.05, \*\**P* < 0.01, \*\*\**P* < 0.001).

## Data availability

The data that support the findings of this study are available on request from the corresponding author.

## Acknowledgments

We thank Dr. Kun-Liang Guan and Dr. Alain Mauviel for providing cDNAs of components of Hippo pathway, and Dr. Hiroshi Sasaki for the 8xGT-IIC-δ51LucII construct. We are grateful to Jiajia He for her excellent technical assistance, and Qingwei Zhu for helpful suggestions, discussions and help with experimental procedures. We also thank Denise Schichnes and Steve Ruzin at the CNR biological imaging facility at the University of California, Berkeley for assistance with microscopy. This study was supported by DOD/U.S. ARMY MEDICAL RESEARCH AND MATERIEL COMMAND W81XWH-15-1-0068 to K.L. and Q.Z., a Tel Aviv University-University of California Berkeley collaborative research grant to Y.I.H. and K.L., and NIH R01AI41757 to Q.Z. Y.I.H. is an incumbent of the Zalman Weinberg Chair in Cell Biology. T.W. is supported by the China Scholarship Council, and Y.L. is supported by the Tang Scholars program.

## Author contributions

T.W., Y.L., and K.L. designed the research. T.W. performed *in vitro* experiments. Y.L. performed *in vivo* experiments. Y.H. designed and O.G. performed FRAP experiments. T.W., Y.L., H.L., Y.H., Q.Z. and K.L. analyzed data and wrote the paper. K.L. conceived and directed the project. All authors discussed the results and commented on the manuscript.

## Supplementary Figure Legends

**Supplementary Fig. 1.**
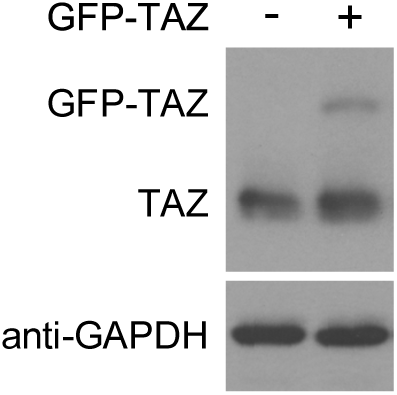
The expression level of GFP-TAZ protein in transfected MCF-10A cells. Ectopically expressed GFP-TAZ was expressed at a lower level than endogenous TAZ in MCF-10A cells as shown by western blotting. GAPDH was used as a loading control.

**Supplementary Fig. 2.**
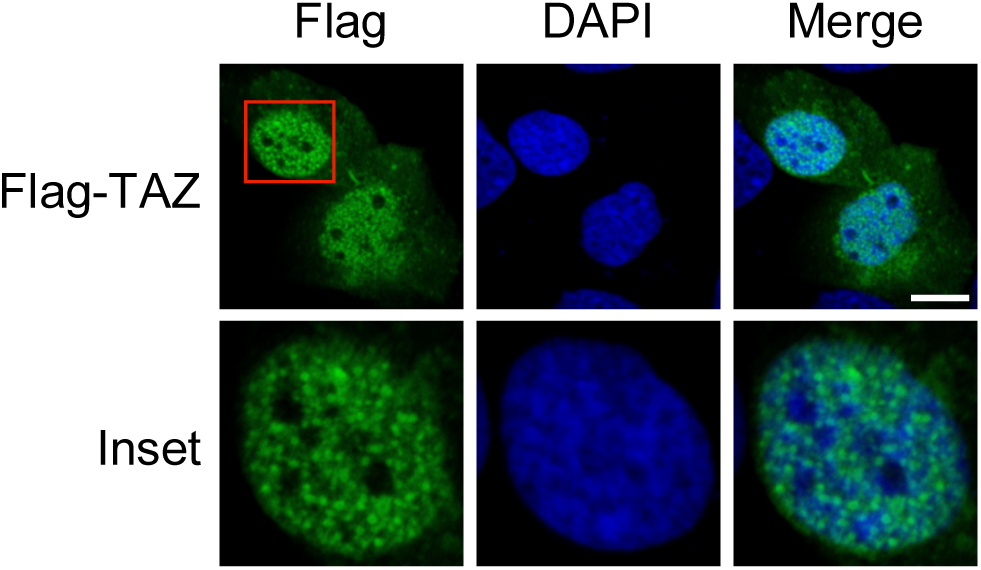
Flag-TAZ forms nuclear puncta in transfected MCF-10A cells. Flag-TAZ formed nuclear puncta when transfected into the MCF-10A cells, as detected by immunofluorescence staining with anti-Flag. Scale bar, 10 μm.

**Supplementary Fig. 3.**
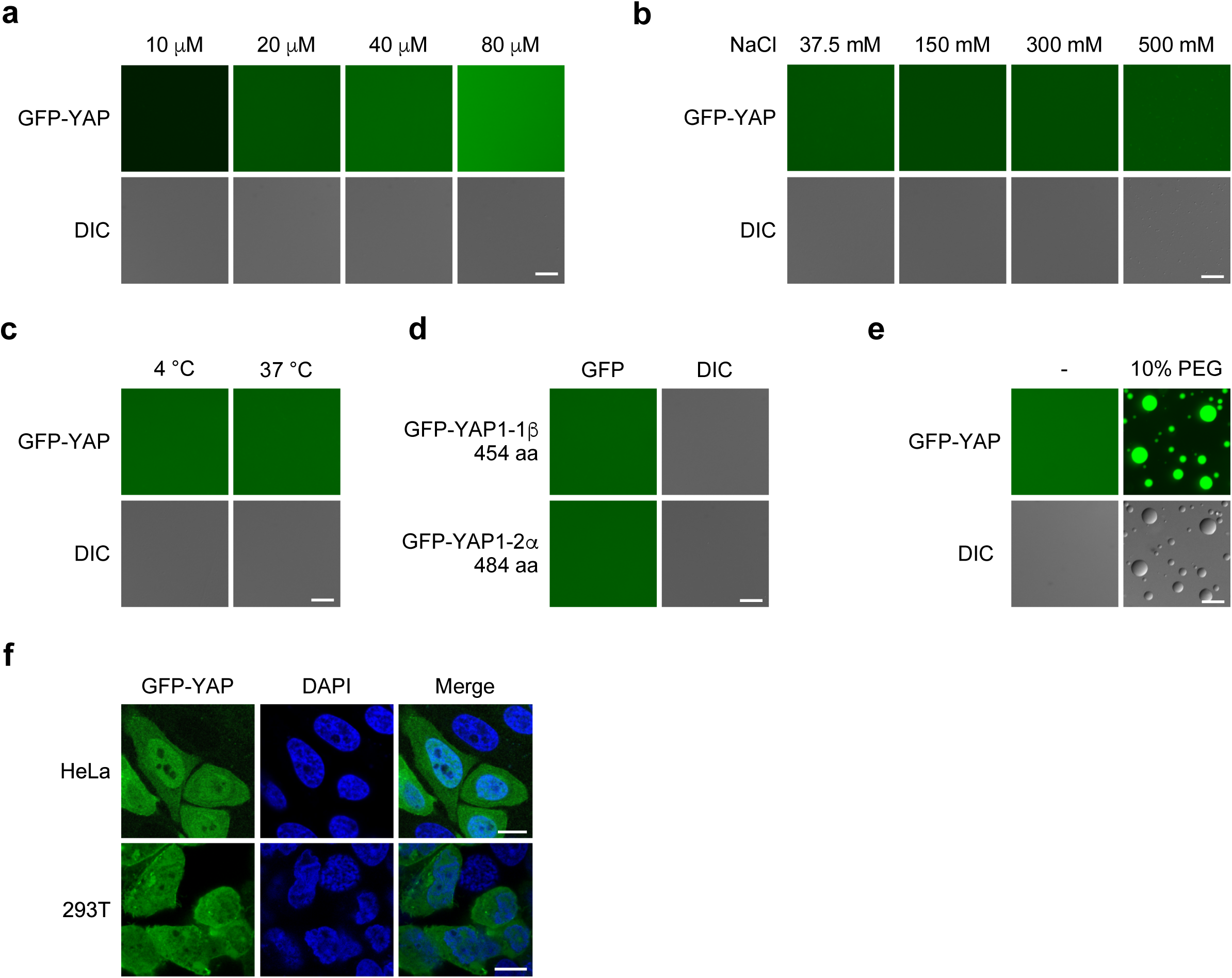
YAP does not form droplets *in vitro* and *in vivo*. **a**) GFP-YAP at varying concentrations was subjected to the droplet formation assay at room temperature and in the presence of 500 mM NaCl. **b**) 50 μM GFP-YAP was subjected to the droplet formation assay at room temperature in the presence of indicated salt concentrations. **c**) 50 μM GFP-YAP was subjected to droplet formation in the presence of 150 mM NaCl at 4°C or 37°C. **d**) Two YAP isoforms, GFP-YAP1-1β or GFP-YAP1-2α, did not form droplets (50 μM protein, 150 mM NaCl and room temperature). aa, amino acids. **e**) 50 μM GFP-YAP could form droplets in the presence of 10% PEG and 500mM NaCl at room temperature. **f**) GFP-YAP did not form nuclear puncta in both HeLa cells and 293T cells. Scale bar for all panels, 10 μm.

**Supplementary Fig. 4.**
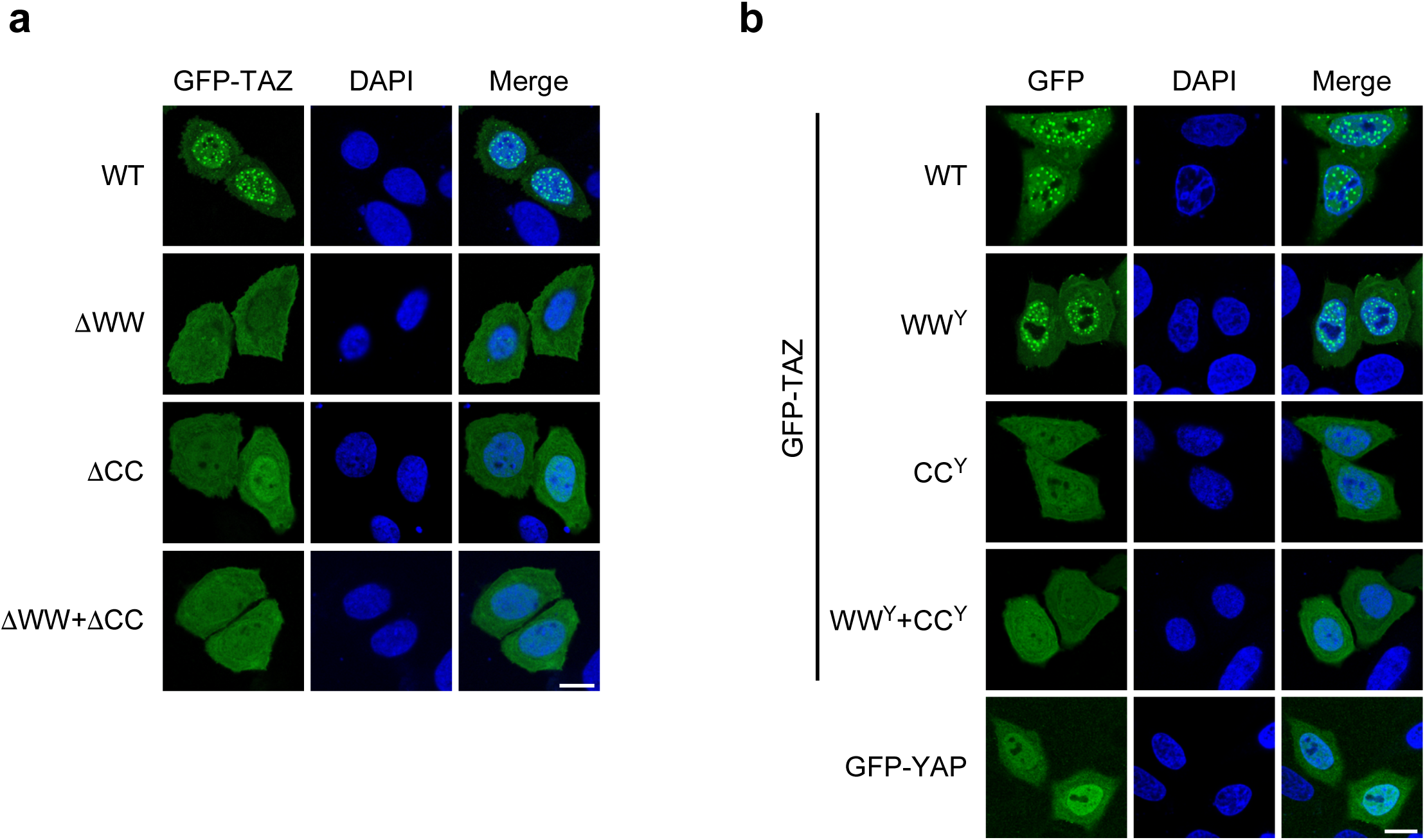
The CC domain is required for TAZ to form nuclear puncta. **a**) Localization of GFP-TAZ and various mutants in HeLa cells. **b**) Localization of GFP-TAZ and various TAZ/YAP chimera in HeLa cells. Scale bar, 10 μm.

**Supplementary Fig. 5.**
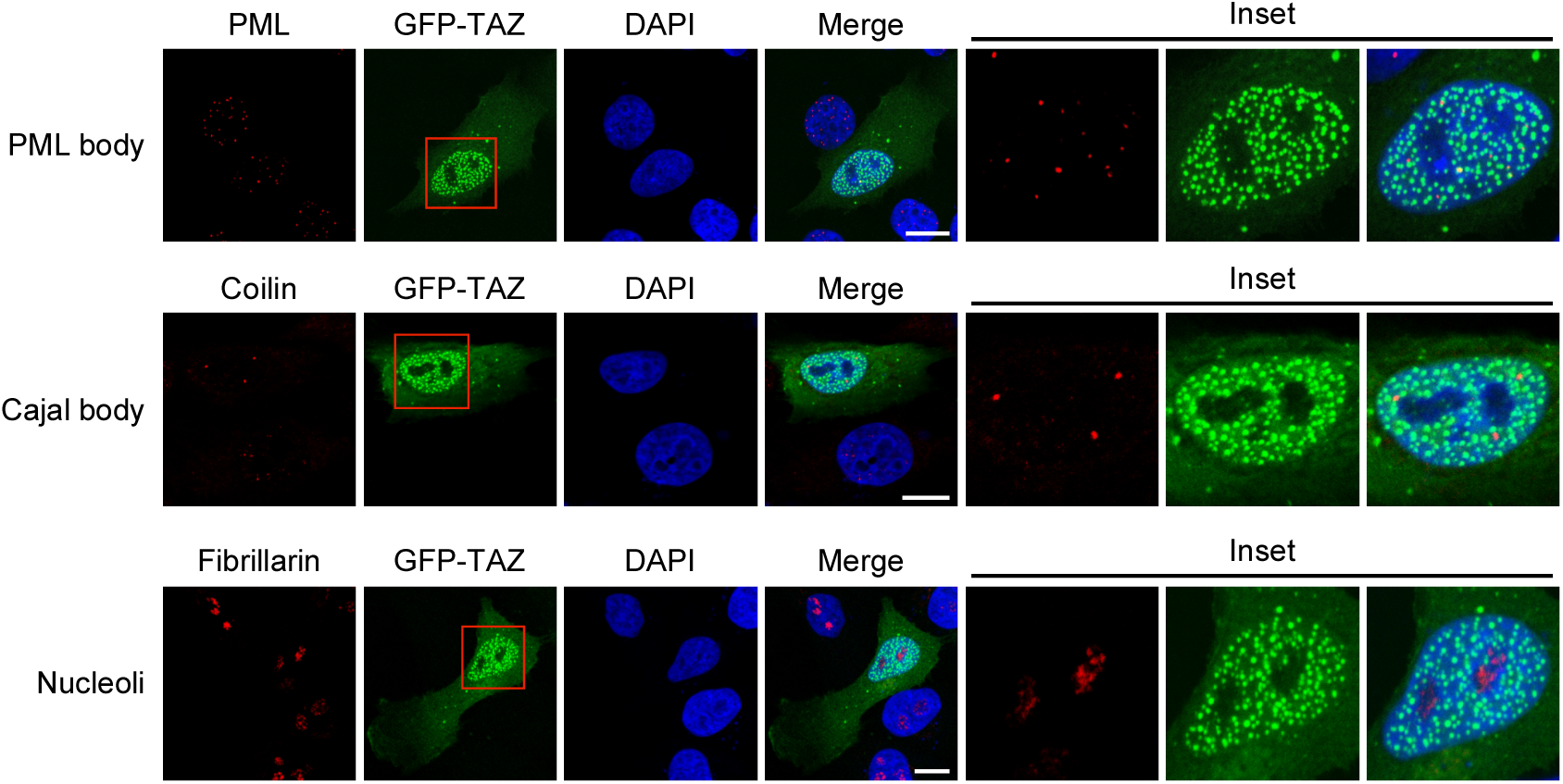
TAZ nuclear condensates do not co-localized with the PML bodies, Cajal bodies or nucleoli. The PML nuclear bodies, Cajal Bodies and nucleoli in MCF-10A cells expressing GFP-TAZ (green) were detected by immunofluorescence staining with antibodies targeting PML, Coilin and Fibrillarin, respectively (red). Scale bar, 10 μm.

**Supplementary Fig. 6.**
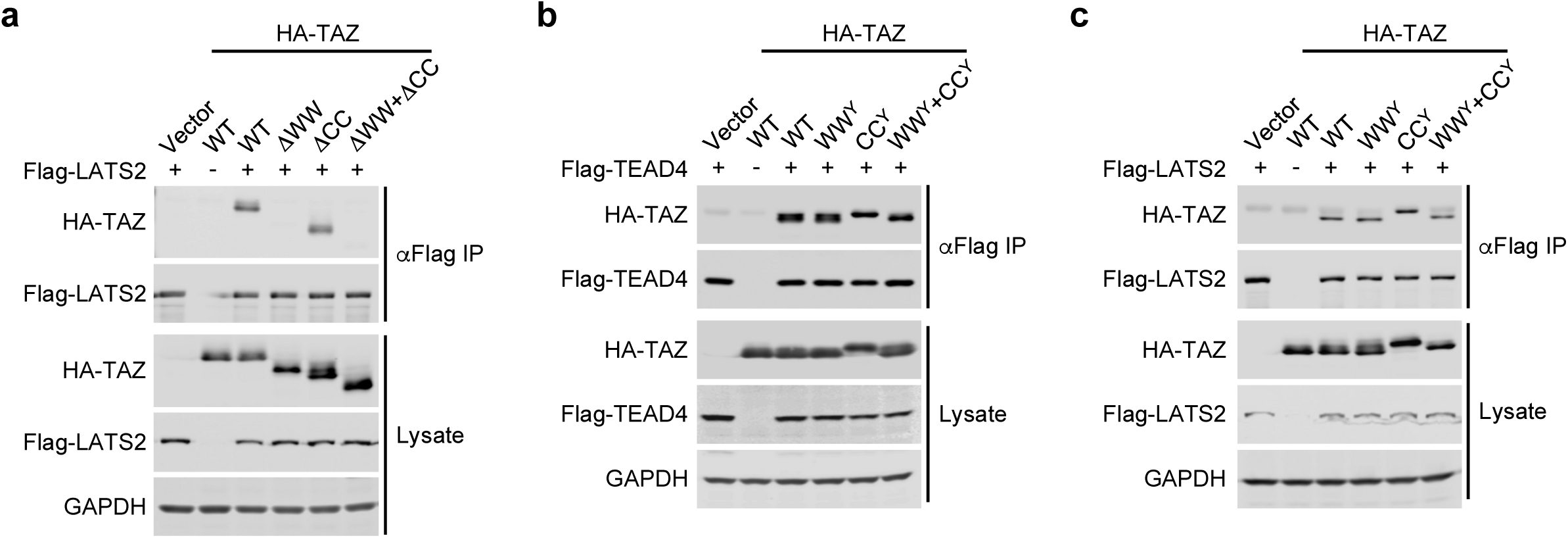
TAZ mutants lacking the CC domain still bind to LAST2 and TEAD4. **a**) HA-tagged WT and mutant TAZ were co-transfected into 293T cells with Flag-LATS2. TAZ proteins associated with LATS2 were isolated by immunoprecipitation with anti-Flag and detected by western blotting with anti-HA antibodies (upper panels). The abundance of these proteins in the cell lysates was assessed by western blotting (lower panels). GAPDH was used as a loading control. **b**). Interaction of various TAZ mutants with Flag-TEAD4 was analyzed by co-IP assay as described in (**a**). **c**) Interaction of various TAZ/YAP chimera with LATS2 was analyzed by co-IP as described in (**a**).

